# Impaired motor activity in a CRISPR SCA5 L253P knock-in mouse is associated with selective β-III-spectrin subcellular redistribution in the cerebellum

**DOI:** 10.64898/2026.03.14.711824

**Authors:** Adam W. Avery, Brennon L. O’Callaghan, Matthew T. Thiel, Sarah A. Denha, Devon G. O’Callaghan, Emma M. Cismas, Jared Lamp, Harry T. Orr, Thomas S. Hays

**Author notes:** Co-first authors (Adam W. Avery, Brennon L. O’Callaghan). Co-senior authors (Harry T. Orr, Thomas S. Hays). Corresponding Authors: Thomas S. Hays, Harry T. Orr and Adam W. Avery.

## Abstract

The spinocerebellar ataxia type 5 (SCA5) L253P mutation in β-III-spectrin causes high-affinity actin binding. Here we developed a CRISPR knock-in mouse to determine the i*n vivo* impact of L253P on Purkinje neurons and motor activity, and to establish a model for future testing of SCA5 therapeutics. Significantly, the knock-in mouse shows impaired motor activity on elevated beam assays at 20 weeks. In the cerebellum, L253P causes a subcellular redistribution of β-III-spectrin in Purkinje neurons. This is marked by loss of β-III-spectrin in distal dendrites, accumulation of β-III-spectrin at the plasma membrane of the soma and proximal dendrites, and formation of inclusions in the soma. The inclusions additionally contain F-actin and α-II-spectrin, accumulate around the nucleus, form at an early age, and are larger in homozygous *β-III-spectrin*^*L253P/L253P*^ compared to heterozygous *β-III-spectrin*^*L253P/+*^ mice. In contrast, neurons of the hippocampus and cerebral cortex, where β-III-spectrin is also known to be expressed, abnormally accumulate β-III-spectrin at the plasma membrane but do not form inclusions. To gain greater insight into disease mechanisms, unbiased proteomics identified over 150 cerebellar proteins that physically associate with β-III-spectrin. Of these, cluster analysis revealed a group of 41 proteins, including glutamate receptors, SERCA2, and CaMKII, linked to synaptic transmission. Thus, the effect of the L253P to alter β-III-spectrin localization, including decreased levels in distal dendrites, is likely associated with a disruption of β-III-spectrin function in postsynaptic signaling. Consistent with this, and in agreement with prior findings in knockout mice, the L253P *β-III-spectrin* knock-in mouse here shows that CaMKII, a calcium sensor and key mediator of glutamate signaling, is ~2-fold activated. Further, the abundance of EAAT4, a glutamate transporter, is significantly reduced. The L253P knock-in mouse primes future preclinical testing of SCA5 therapeutics, such as small molecule modulators of spectrin-actin binding, and glutamate and calcium signaling pathways.

## Introduction

Spinocerebellar ataxia type 5 (SCA5) is an autosomal dominant neurodegenerative disorder caused by mutations in the *SPTBN2* gene encoding β-III-spectrin (1). One of the first SCA5 mutations identified was a missense mutation resulting in substitution of leucine 253 for proline (L253P) (1). The pathological consequences of L253P were characterized in fifteen members of a German family (2). Symptoms included ataxia of gait, limb and stance, dysarthria, and downbeat nystagmus. Symptoms began at an average age of 32 years and showed slow progression. MRI revealed cortical cerebellar atrophy but sparing of other brain regions (2). Consistent with targeting of the cerebellum, β-III-spectrin is known to be highly expressed in cerebellar Purkinje neurons (3). Molecularly, β-III-spectrin contains an N-terminal actin-binding domain (ABD), seventeen spectrin-repeat domains (SRDs) and a C-terminal pleckstrin homology domain. Significantly, L253P localizes to the β-III-spectrin ABD and increases actin-binding affinity by approximately 1000-fold (4, 5). Recently, many additional ABD-localized SCA5 mutations were shown to increase actin-binding affinity (6), supporting increased actin binding as a common SCA5 molecular consequence. Indeed, the identification of small molecules that modulate the binding of L253P β-III-spectrin to actin is being pursued as a possible SCA5 therapeutic (7, 8). However, an appropriate in vivo model for L253P has yet to be developed for testing of potential SCA5 therapeutics. Such a model is additionally needed to gain greater mechanistic insights into how increased actin binding disrupts Purkinje neuron function and motor activity.

It is known that β-III-spectrin binds α-II-spectrin to form a heterotetramer (9–11). The heterotetramer crosslinks actin filaments via the β-III-spectrin ABDs, positioned at opposite ends of the tetramer. This results in a spectrin-actin cytoskeleton that lines the plasma membrane and is present throughout the somatodendritic compartment of Purkinje neurons, and of neurons in the cerebral cortex and hippocampus (9, 10, 12, 13). Mouse studies using dissociated Purkinje neurons transfected *in vitro*, or electroporated immature Purkinje neurons in utero, showed that the ectopically expressed L253P mutant is present in the soma, but forms “aggregates.” Further the L253P mutant protein showed dramatically reduced localization to the dendritic arbor (10). These transfected Purkinje neurons further showed a loss of dendritic arbor planarity. Earlier studies in Drosophila dendritic arborization (da) neurons similarly showed that overexpressed fly β-spectrin containing the L253P mutation accumulates in the cell body and proximal dendrites but is absent in distal dendrites (14). In the L253P da neurons, this proximal shift in subcellular localization of the mutant was associated with distal dendrite degeneration. Proximal dendrites continue to grow, but the outward growth of the arbor is impaired. Altogether, these studies in electroporated/transfected Purkinje neurons and Drosophila da neurons established that high-affinity actin binding is associated with a subcellular redistribution of β-III-spectrin, and that the absence of β-III-spectrin in dendrites disrupts dendrite stability and growth. However, whether endogenous expression of a knock-in mutant allele causes similar β-III-spectrin redistribution and the impact of the redistribution on Purkinje neuron function are not known.

Information on β-III-spectrin and its importance in the cerebellum comes from two independently developed knockout mouse models. These mice showed that loss of β-III-spectrin causes severe, early motor impairment and cerebellar atrophy (12, 15). In these mice, Purkinje neurons show decreased number of dendritic spines, reduced dendritic arbor planarity, and progressive arbor degeneration (12, 15, 16). Further, loss of β-III-spectrin causes multiple synaptic proteins including the glutamate transporter, EAAT4, to be decreased in abundance (12, 15, 16). The decrease in EAAT4 was suggested to contribute to excessive synaptic glutamate signalling, leading to Purkinje neuron excitotoxicity and ataxia in the knockout mouse (15, 17–19). In agreement with this model, *EAAT4* knockout mice show progressive motor impairment (18). Further, in the *β-III-spectrin* knockout mouse, intracellular calcium signaling is elevated, as monitored by an increased activation state of CaMKII (19), supportive of excessive glutamate excitation. Thus, *in vivo* studies are needed to determine whether dominant SCA5 mutations, such as L253P, disrupt calcium signalling and EAAT4 abundance.

A transgenic mouse model was previously generated to examine the impact of a SCA5 in-frame deletion, E532_M544del, localized to spectrin-repeat domain 3. In this model, transgenic wild-type or mutant human β-III-spectrin was conditionally expressed in Purkinje neurons (20). In contrast to the knockout mice, the E532_M544del mouse showed mild motor impairment on the rotarod, and mild cerebellum atrophy at late time points. Significantly, the E532_M544del mutant did not disrupt EAAT4 protein level, consistent with normal EAAT4 protein levels measured in autopsy tissue from a patient with the same mutation (1). Further, the glutamate receptor, mGluR1α, was identified as a β-III-spectrin interacting protein (20). E532_M544del disrupted the organization of mGluR1α at post-synaptic sites, which was associated with diminished mGluR1α signaling. Thus, the E532_M544del mouse study suggests that pathogenesis induced by dominant SCA5 mutations may be distinct from the degenerative mechanisms triggered by loss of β-III-spectrin in knockout mice. Further, this work highlights the need to more fully understand the protein interactions of β-III-spectrin in cerebellum, to improve our understanding of β-III-spectrin function and the molecular pathways impacted by SCA5 mutations. In the present study, CRISPR-Cas9 engineering was performed to introduce the L253P mutation in the mouse *Sptbn2* gene, resulting in a SCA5 knock-in mouse model. The mouse shows mild motor impairment that is associated with the subcellular redistribution of β-III-spectrin from distal dendrites to proximal dendrites and the soma. We show that the mutant β-III-spectrin forms inclusions in Purkinje neurons, but not hippocampal or cortical neurons. These inclusions additionally contain α-II-spectrin and F-actin and coalesce around the nucleus. Unbiased proteomics using cerebellar tissue identified many known and novel β-III-spectrin-associated proteins, including CaMKII and SERCA2, highlighting a role for β-III-spectrin in post-synaptic signaling. Consistent with this role, and with prior findings in knockout mice, in the L253P mouse here, we find that CaMKII activity is elevated and EAAT4 protein level is reduced. This work establishes an important SCA5 in vivo model for understanding how increased actin binding disrupts Purkinje neuron function and motor activity, and for future testing of SCA5 therapeutics.

## Results

### Development of the CRISPR L253P knock-in mouse

To characterize the in vivo impact of L253P, CRISPR-Cas9 technology was used to introduce the L253P mutation into the endogenous *Sptbn2* gene through homology-directed repair. The repair template introduced a single base change to convert the leucine codon “CTC” to a proline codon “CCC”, **Fig. 1A**. Additionally, a silent mutation was introduced that converted the “TGG” PAM, located adjacent to the gRNA target sequence, to “TAG”, to minimize unwanted extraneous non-homologous end joining. Further the PAM mutation introduced a BfaI restriction site (CTAG) to aid in screening of successful homology-directed repair. Sanger sequencing confirmed successful genome editing, **Fig. 1B**, and a stable breeding population containing the L253P mutation was established.

**Fig. 1.**
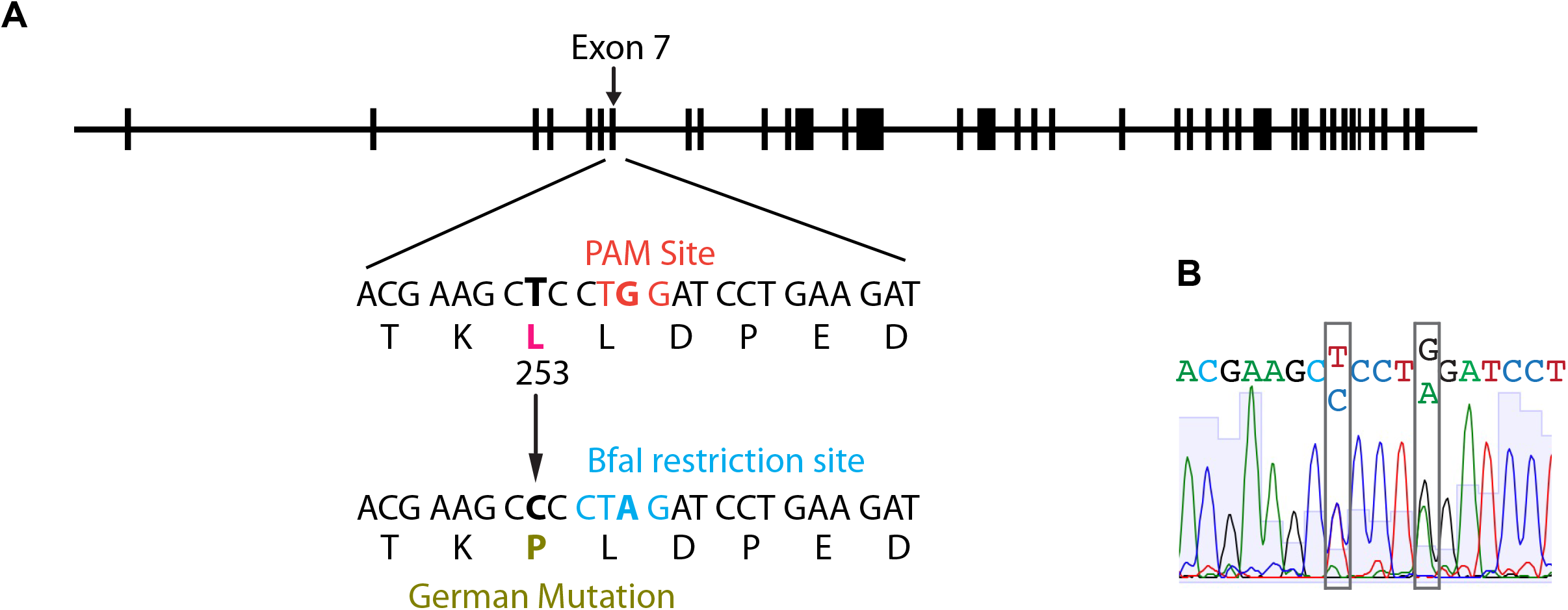
CRISPR-Cas9 introduction of the L253P mutation. (A) The L253P (German) mutation was introduced into the *Sptbn2* gene via CRISPR-Cas9 homology-directed repair. The PAM site was destroyed with a silent mutation that simultaneously introduced a BfaI restriction site. (B) Sanger sequencing confirmed the heterozygous mutation.

To determine the impact of L253P on β-III-spectrin expression, RT-qPCR was performed on cerebellum from wild-type (WT/*β-III-spectrin*^*+/+*^), heterozygous *β-III-spectrin*^*L253P/+*^ and homozygous *β-III-spectrin*^*L253P/L253P*^ mice, **Fig. 2A**. No difference in transcript level was observed between WT and *β-III-spectrin*^*L253P/+*^ mice. However, a ~20% increase in transcript level was observed in *β-III-spectrin*^*L253P/L253P*^ mice relative to WT, suggesting modest compensatory transcriptional upregulation of the Sptbn2 gene. To assess β-III-spectrin protein expression, cerebellar extracts were prepared from WT, *β-III-spectrin*^*L253P/+*^ and *β-III-spectrin*^*L253P/L253P*^ animals, and equal amounts of total protein were analyzed by SDS-PAGE followed by western blot with β-III-spectrin antibody, **Fig. 2B**. In the detergent (Triton X-100 and IGEPAL) soluble fractions, β-III-spectrin protein level was decreased by ~50% in β-III-spectrin^L253P/+^ animals and by nearly 100% in *β-III-spectrin*^*L253P/L253P*^ animals, relative to WT, **Fig. 2C**. Heat and SDS were used to extract additional β-III-spectrin contained in the cerebellar insoluble material. β-III-spectrin protein was successfully extracted from the insoluble fraction of *β-III-spectrin*^*L253P/L253P*^ tissue, although less was recovered than the amount extracted from WT or *β-III-spectrin*^*L253P/+*^ tissue, **Fig. 2D**. Altogether, these data show L253P β-III-spectrin is expressed at the mRNA and protein level. However, L253P shifts β-III-spectrin from the soluble to the detergent insoluble cerebellar fraction.

**Fig. 2.**
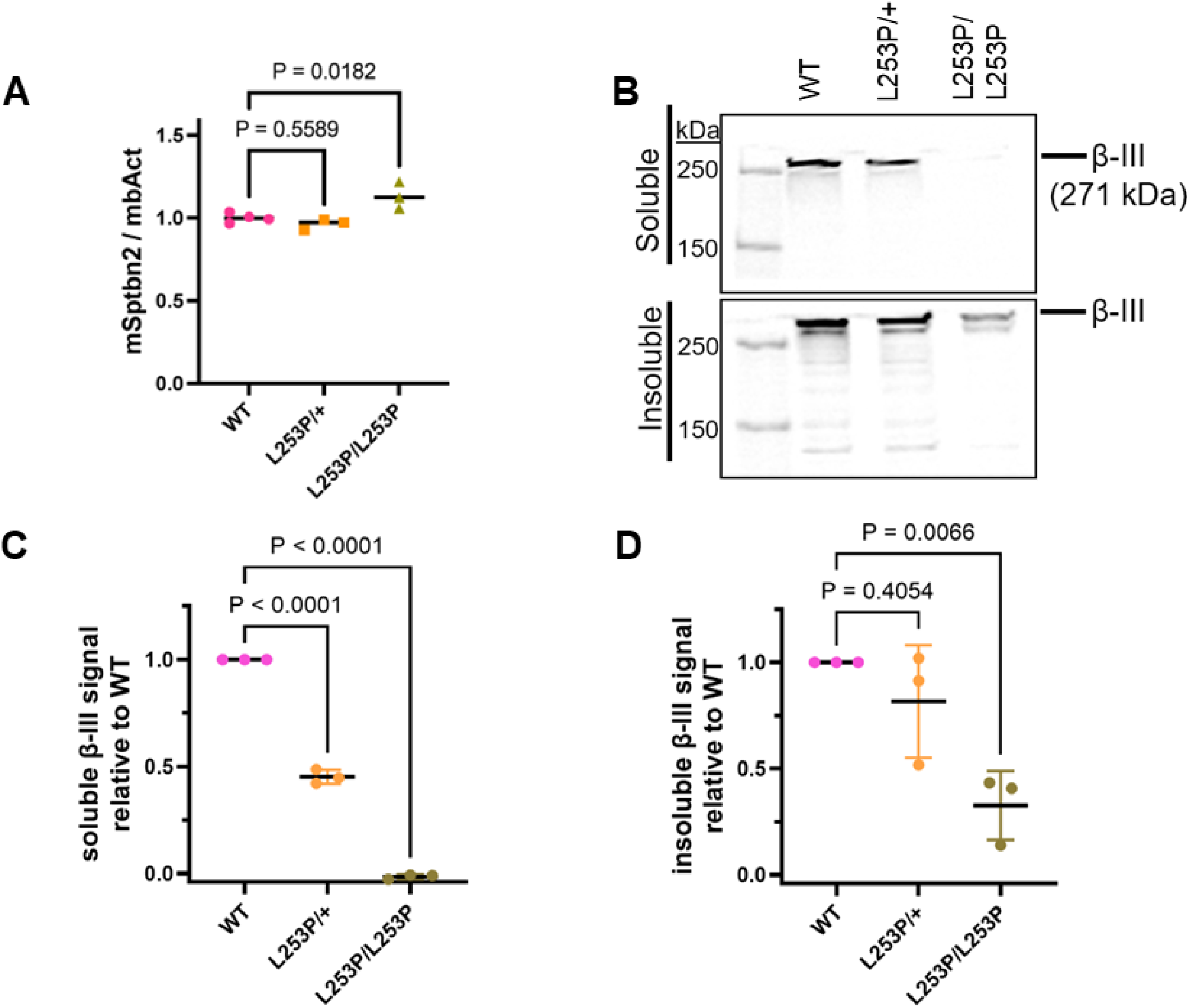
Expression of β-III-spectrin in mouse cerebellum. (A) RT-qPCR shows a modest ~20% increase in homozygous β-III-spectrin transcript levels, n=3-4 (B-D) Western blot of cerebellum extracts shows progressive decrease in β-III-spectrin protein in heterozygous and homozygous mice in both soluble and insoluble fractions, n=3. (A, C, D) One way ANOVA comparing to wild-type using Dunnett’s multiple comparison correction.

### L253P causes mild impairment of motor behaviour

To determine how L253P impacts motor behaviour, mice were initially tested on the rotarod at 6 and 24 weeks. However, no differences in motor performance were observed across the genotypes, at either age, **Fig. S1**. Subsequently, mice were assayed on an elevated beam for foot slips. Beams of different diameter (10 mm versus 16 mm) and shape (round versus square) were tested. On the elevated beam, young (six weeks) mice of any genotype rarely displayed foot slips on any beam type, **Fig. 3**. In contrast, at 20 weeks, *β-III-spectrin*^*L253P/L253P*^ mice showed a statistically significant increase in number of foot slips on all beam types tested. Notably, on the round, 10 mm beam, the most challenging for mice to cross, *β-III-spectrin*^*L253P/+*^ animals showed an increase in foot slips that approached the threshold for statistical significance (p = 0.053). Thus, *β-III-spectrin*^*L253P/L253P*^ animals display reproducible motor impairment by 20 weeks. Moreover, at 20 weeks, *β-III-spectrin*^*L253P/+*^ animals are also likely impaired, but to a lesser extent than *β-III-spectrin*^*L253P/L253P*^.

**Fig. 3.**
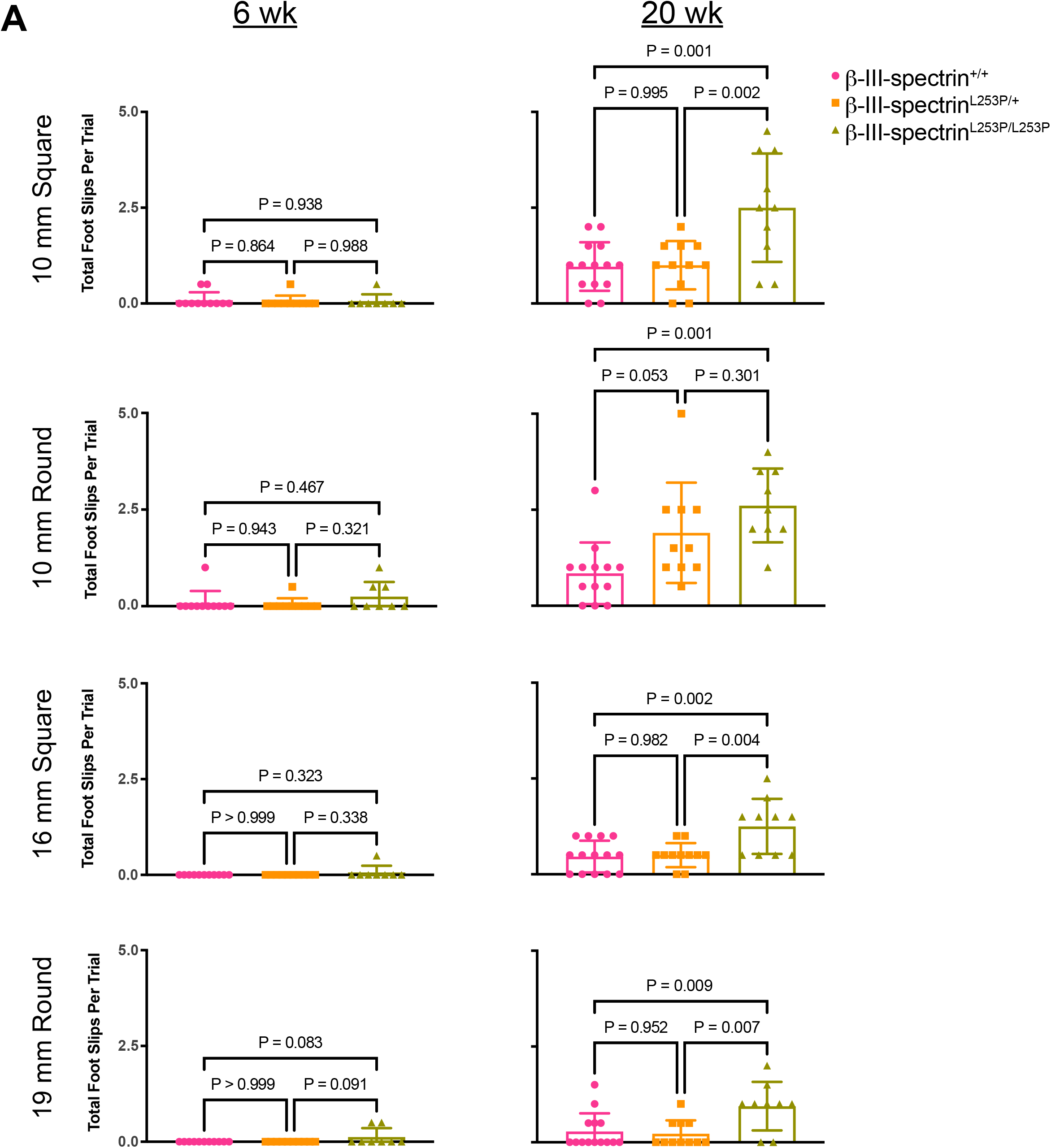
L253P causes an age dependent loss of motor coordination. (A) Number of foot slips 6 week and 20 week old mice on 10 or 16 mm, square or round elevated beams. At 6 weeks, no increase in foot slips is observed on any beam. At 20 weeks, *β-III-spectrin*^*L253P/L253P*^ mice show increased foot slips on the 10 and 16 mm round and square beams. *β-III-spectrin*^*L253P/+*^ mice show increased number of foot slips that approaches statistical significance on the 10 mm round beam. One way ANOVA followed by multiple comparisons with Tukey’s correction, n=10-14

### L253P redistributes β-III-spectrin from the Purkinje neuron dendritic arbor to the soma

To determine how L253P impacts the subcellular localization of β-III-spectrin, immunofluorescence was performed on cerebellum slices from 20-week mice. In WT mice, using an antibody targeting the C-terminus of β-III-spectrin, β-III-spectrin appears as a diffuse signal in the soma and throughout the dendritic arbor, counter-labelled using an antibody against calbindin, **Fig. 4A**. In contrast, *β-III-spectrin*^*L253P/L253P*^ mice show a marked loss of β-III-spectrin signal within the dendritic arbor. Instead, β-III-spectrin becomes concentrated within the soma, where it appears as prominent, punctate structures. Although these structures were previously described as “aggregates” (10), this term implies an uncharacterized biophysical state; therefore, we refer to them here as “inclusions.” This redistribution was confirmed using an independent antibody targeting the N-terminus of β-III-spectrin, **Fig. S2**. A closer examination of the soma and proximal dendrites, across the three genotypes, revealed additional information. In contrast to the diffuse soma β-III-spectrin signal observed in Purkinje neurons of WT animals, in *β-III-spectrin*^*L253P/+*^ animals, β-III-spectrin forms small inclusions in the soma, and accumulates at the plasma membrane, such that the soma and proximal dendrites appear outlined, **Fig. 4B**. In *β-III-spectrin*^*L253P/L253P*^ tissue, β-III-spectrin staining shows the plasma membrane of the soma and proximal dendrites to be much more intense and a mix of small and very large inclusions are present. This subcellular redistribution of β-III-spectrin is apparent from an early age, with inclusions occupying ~25% of the volume of the soma in *β-III-spectrin*^*L253P/L253P*^ mice, measured at 6 weeks and 20 weeks, **Fig. 4C**. Together these findings show that the L253P mutation drives a pronounced shift in β-III-spectrin localization away from distal dendritic arbor toward the plasma membrane of the soma and proximal dendrites, accompanied by inclusion formations within the soma. The more pronounced redistribution of β-III-spectrin in *β-III-spectrin*^*L253P/L253P*^ animals compared to *β-III-spectrin*^*L253P/+*^ is consistent with a dosage dependent effect of the mutant.

**Fig. 4.**
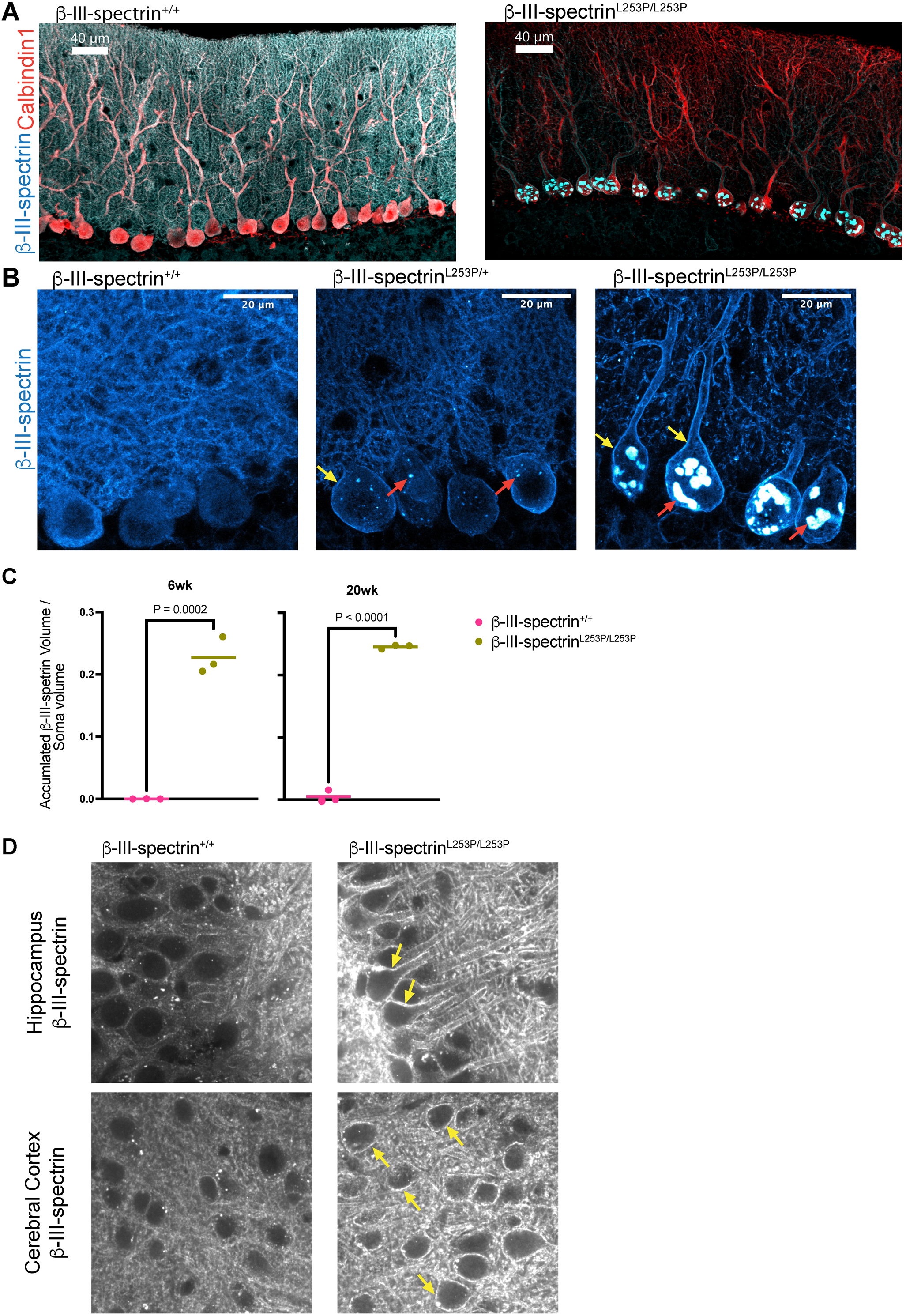
L253P causes a pronounced subcellular redistribution of β-III-spectrin. (A) Immunofluorescent labelling of β-III-spectrin shows distribution changes in the soma and throughout the dendritic arbor of 20 week mice. (B) Close-up view of soma and proximal dendrites. In WT mice, β-III-spectrin shows a diffuse distribution in the soma and proximal dendrites. In *β-III-spectrin*^*L253P/+*^ mice, β-III-spectrin shows accumulation at the plasma membrane of the soma and proximal dendrites (yellow arrows) and forms small inclusions in the soma (red arrows). In *β-III-spectrin*^*L253P/L253P*^ mice, β-III-spectrin localizes strongly to the proximal dendrites and soma plasma membrane with small and large intracellular inclusions present. (C) Quantitation of the total inclusion volume relative to the soma volume in 6 week and 20 week wild-type and homozygous mice. Unpaired t-test, n=3 mice per group with at least 10 somas per image. (D) Hippocampal pyramidal neurons and cortical neurons showing plasma membrane accumulation (yellow arrows) of β-III-spectrin in *β-III-spectrin*^*L253P/L253P*^ mice, but no inclusions.

L253P is thought to only impact the cerebellum (2). However, β-III-spectrin is known to be expressed in other brain regions including the hippocampus and cerebral cortex (12, 13). Thus, we examined the subcellular localization of β-III-spectrin in hippocampal and cortical neurons. In hippocampal pyramidal neurons of WT mice, β-III-spectrin appears primarily as a diffuse signal in the soma and dendrites, with only faint, occasional plasma membrane labeling, **Fig. 4D**. However, in *β-III-spectrin*^*L253P/L253P*^ mice, β-III-spectrin shows pronounced plasma membrane localization, strongly outlining the soma and dendrites. In contrast to Purkinje neurons, in the pyramidal neurons, no inclusions are detected. Similarly, in cortical neurons of *β-III-spectrin*^*L253P/L253P*^ mice, β-III-spectrin accumulates at the cell cortex but does not form inclusions. Thus, inclusions, a prominent feature of cerebellar Purkinje neurons expressing the L253P mutant, are absent in the neurons of other brain regions known to express β-III-spectrin. Given that L253P selectively targets the cerebellum (2), it is possible that the inclusions are a driver of Purkinje neuron dysfunction or reflect L253P disruption of a unique pathway critical to Purkinje neuron function.

### L253P β-III-spectrin inclusions contain α-II-spectrin and F-actin and accumulate around the nucleus

Inclusions are a prominent feature of Purkinje neurons expressing the L253P mutant. Thus, we further characterized the inclusions to gain a better understanding of their composition and distribution. Intriguingly, in *β-III-spectrin*^*L253P/L253P*^ tissue, large inclusions are closely associated with the cytoplasmic side of the nuclear envelope, **Fig. 5A**. In contrast, small inclusions are more scattered in the soma or localize near the plasma membrane, **Fig. S3**. Further, the inclusions show strong co-localization with F-actin, **Fig. 5B**. This agrees with the known impact of L253P to strongly increase β-III-spectrin actin binding in vitro (4). In addition, α-II-spectrin, a known high-affinity interactor (11), shows pronounced co-localization with the inclusions, **Fig. 5C**. The presence of F-actin and α-II-spectrin in the inclusions suggests that β-III-spectrin in the inclusions maintains the ability to interact with its naturally occurring binding partners. In contrast, we did not observe EAAT4 (21) or ankyrin-R (22, 23) in the inclusions (**Fig. S4**), supporting that only a subset of β-III-spectrin interactors are contained in the inclusions.

**Fig. 5.**
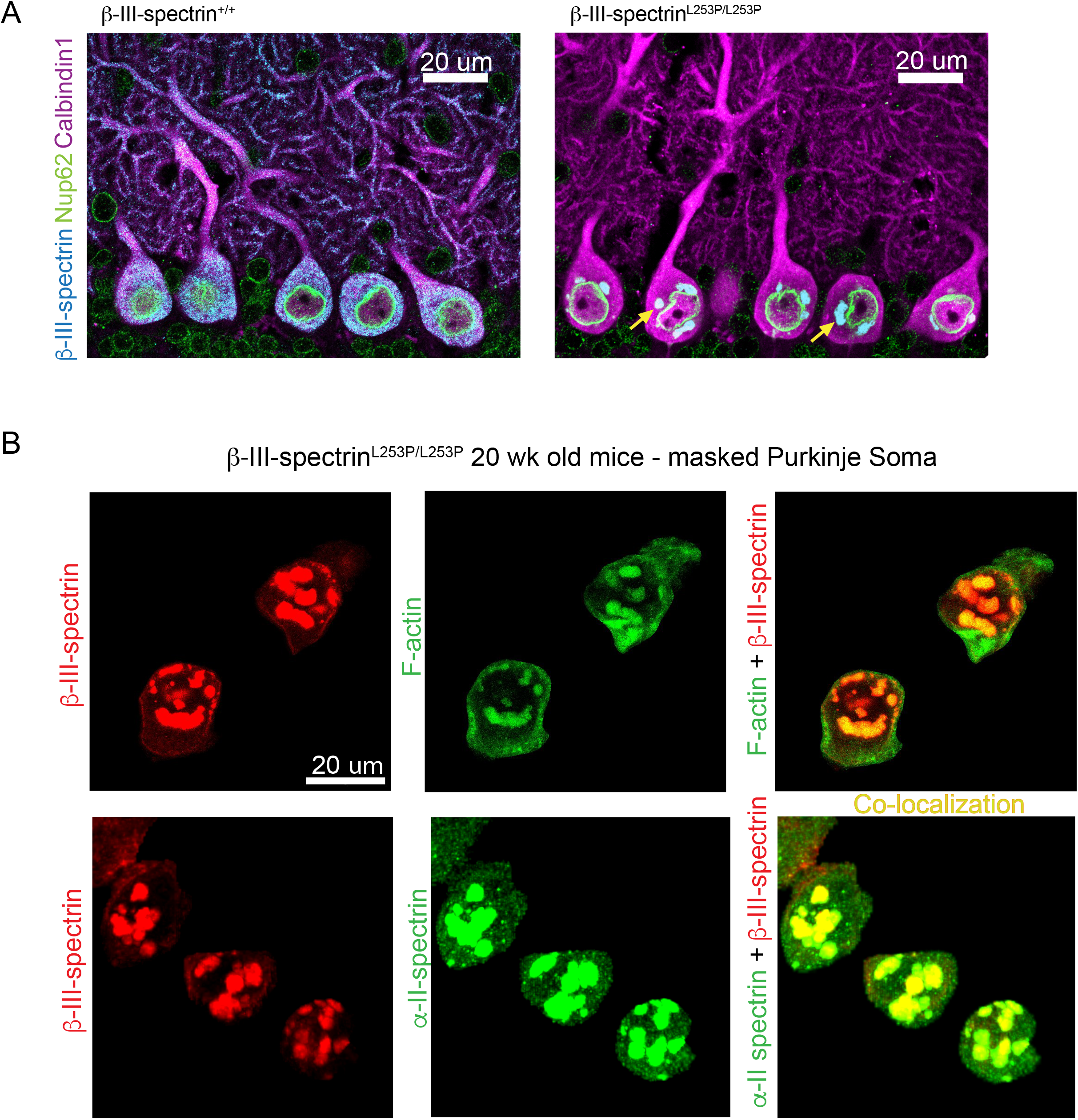
L253P β-III-spectrin inclusions accumulate around the nucleus and contain F-actin and α-II-spectrin. (A) Immunofluorescent labelling of cerebellar sections with nucleoporin 62, β-III-spectrin, and calbindin antibodies shows that inclusions in the *β-III-spectrin*^*L253P/L253P*^ tissue localize around the nucleus, in 20 week mice. (B-C) Calbindin labelled Purkinje neurons somas were masked using Imaris software. (B) Labelling with β-III-spectrin antibody and Alexa-488 phalloidin shows F-actin colocalization with the inclusions. (C) Labelling with β-III-spectrin and α-II-spectrin shows α-II-spectrin colocalization with the inclusions.

### β-III-spectrin physically associates with numerous postsynaptic proteins

To gain a greater understanding of β-III-spectrin physical interactions and associated molecular pathways, immunoprecipitation coupled to tandem mass spectrometry (IP-MS) was performed for unbiased protein identification, using cerebellar tissue extracts. Using WT tissue, β-III-spectrin was enriched by an average of 21.5-fold in the β-III-spectrin antibody IP-MS samples relative to a control IP-MS samples using a non-specific antibody. Thus, as an initial selection criterion, β-III-spectrin interactors were defined as those proteins reproducibly enriched, across triplicate IP-MS experiments, using WT tissue. In **Fig. 2B**, we showed that the amount of soluble β-III-spectrin in *β-III-spectrin*^*L253P/L253P*^ tissue extracts is virtually undetectable by western blot. Consistent with this, in IP-MS experiments using *β-III-spectrin*^*L253P/L253P*^ tissue extracts, relatively very little β-III-spectrin was detected and β-III-spectrin was far less enriched (~1.6-fold). We reasoned that β-III-spectrin interactors would likewise be less abundant and less enriched in *β-III-spectrin*^*L253P/L253P*^ IP-MS samples. Thus, additional selection criteria were that β-III-spectrin interactors must show reduced abundance and reduced enrichment in *β-III-spectrin*^*L253P/L253P*^ IP-MS samples. Meeting these criteria were many known β-III-spectrin interactors, including α-II-spectrin (11), ankyrin-R (22, 23), and mGluR1(20), validating our approach to identify β-III-spectrin associated proteins.

In total the interactome contains 157 proteins that likely directly or indirectly physically associate with β-III-spectrin in the cerebellum, **Fig. S5**. Cluster analyses performed using the STRING database (24) showed that the proteins contained in the interactome are highly connected based on known physical or functional interactions. Numerous clusters were identified, including those related to “chemical transmission across synapse” (41 proteins), “myosin complex” (25 proteins), “cytoplasmic translation (20 proteins), and “COPI-independent Golgi-to-ER retrograde traffic” (13 proteins). **Fig. 6A** displays the large 41-protein cluster associated with synaptic transmission. This cluster contains β-III-spectrin (*Sptbn2*), and many known post-synaptic proteins involved in glutamate and calcium signalling, including the SCA44 disease protein, mGluR1 (*Grm1*) (25), the SCA14 disease protein, protein kinase C gamma (*Prkcg*) (26), the GluN1 subunit the glutamate/glycine activated NMDAR1 calcium channel (*Grin1*), the sarcoplasmic/endoplasmic reticulum calcium ATPase 2 (SERCA2/*Atp2a2*) calcium pump, the calcium/calmodulin dependent serine protein kinase (*Cask*), and components of the CaMKII complex (*Camk2a, Camk2b, Camk2d*). **Fig. 6B** shows several examples of interactor enrichment based on normalized abundance values in the β-III-spectrin antibody versus control antibody IP-MS samples. The high representation in the interactome of proteins involved in synaptic transmission is consistent with the known localization of β-III-spectrin to dendritic membranes and spines (10, 13, 16, 20), and the known requirement of β-III-spectrin to support proper Purkinje neuron post-synaptic signalling (15, 19, 20). Significantly, the impact of L253P to greatly decrease the localization of β-III-spectrin to the dendritic arbor, indicates that the function of β-III-spectrin to regulate of post-synaptic signalling in Purkinje neurons is likely disrupted in L253P mice.

**Fig. 6.**
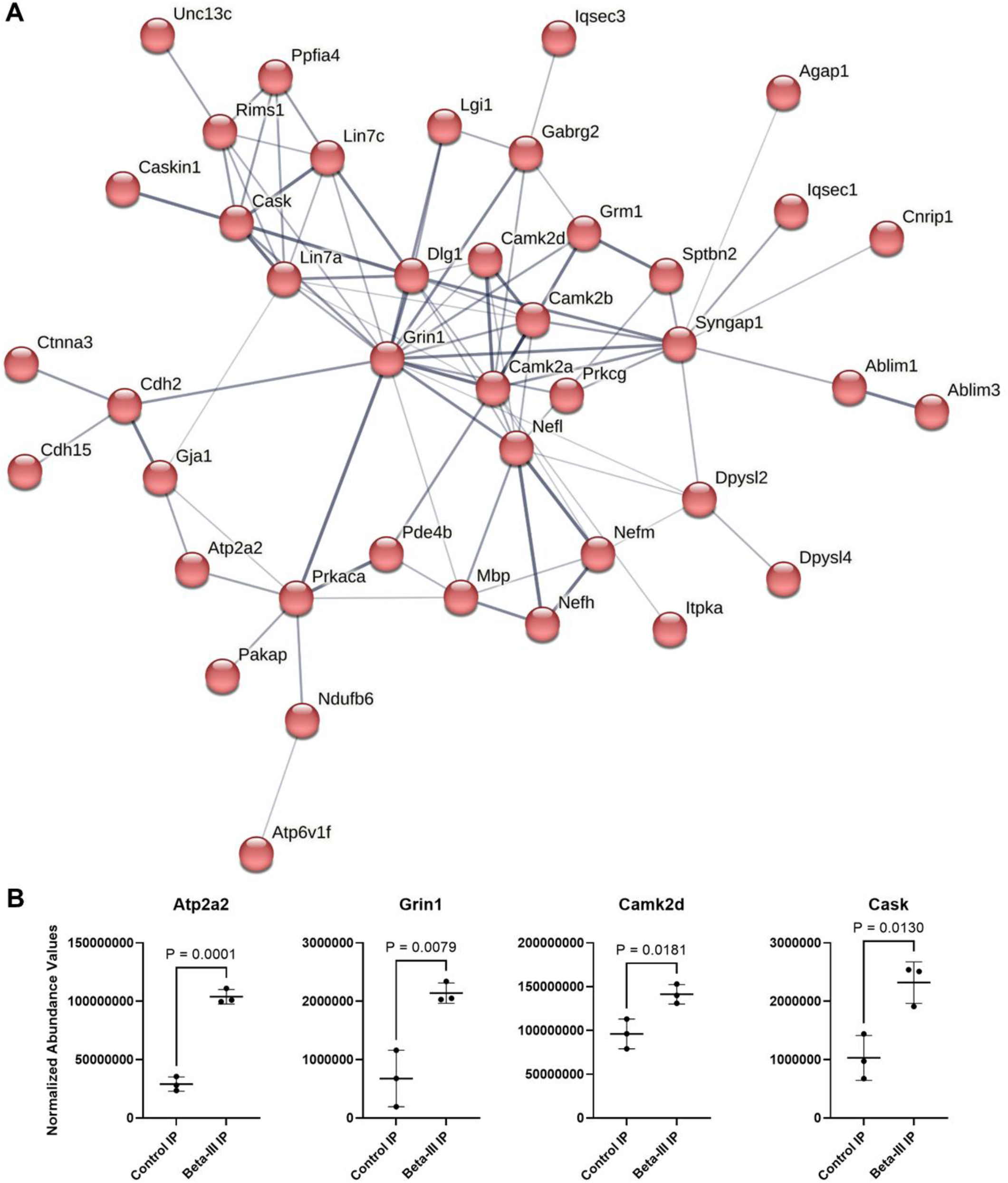
Large cluster of postsynaptic proteins associated with β-III-spectrin in the cerebellum. (A) Immunoprecipitation coupled to tandem mass spectrometry was performed for unbiased identification of proteins that associate with wild-type β-III-spectrin in the cerebellum. STRING identified a cluster of 41 proteins (of 157 total) that are linked to synaptic transmission. Connecting lines represent physical and / or functional linkages between specific proteins, with thicker lines indicating higher confidence interactions. (B) Example proteins from the cluster, with various levels of enrichment, based on normalization abundance values in control versus β-III-spectrin samples. Unpaired t-test, n=3.

### L253P increases CaMKII activation state and reduces EAAT4 abundance

A recent study in *β-III-spectrin* knockout mice showed that the calcium sensor, CaMKII, is activated, based on increased CaMKII auto-phosphorylation and enhanced phosphorylation of known CaMKII targets in the cerebellum (19). This increase in CaMKII activity is an indicator of altered post-synaptic calcium homeostasis. Several subunits of the CaMKII complex and additional regulators of calcium homeostasis, including SERCA2/*Atp2a2*, mGluR1/*Grm1* and GluN1/*Grin1*, are contained in the β-III-spectrin interactome. Together, with the impact of the L253P to reduce β-III-spectrin localization to the dendritic arbor, we reasoned calcium homeostasis may be disrupted in L253P mice. To test this, cerebellar extracts from 25-week mice were probed with an antibody recognizing a specific autophosphorylation site on CaMKII that represents its activation state. Significantly, CaMKII autophosphorylation was increased ~2-fold in *β-III-spectrin*^*L253P/L253P*^ cerebellum extracts compared to WT, **Fig 7A**. Thus, L253P causes activation of the calcium sensor CaMKII, suggesting disrupted calcium homeostasis in the *β-III-spectrin*^*L253P/L253P*^ mouse.

**Fig. 7.**
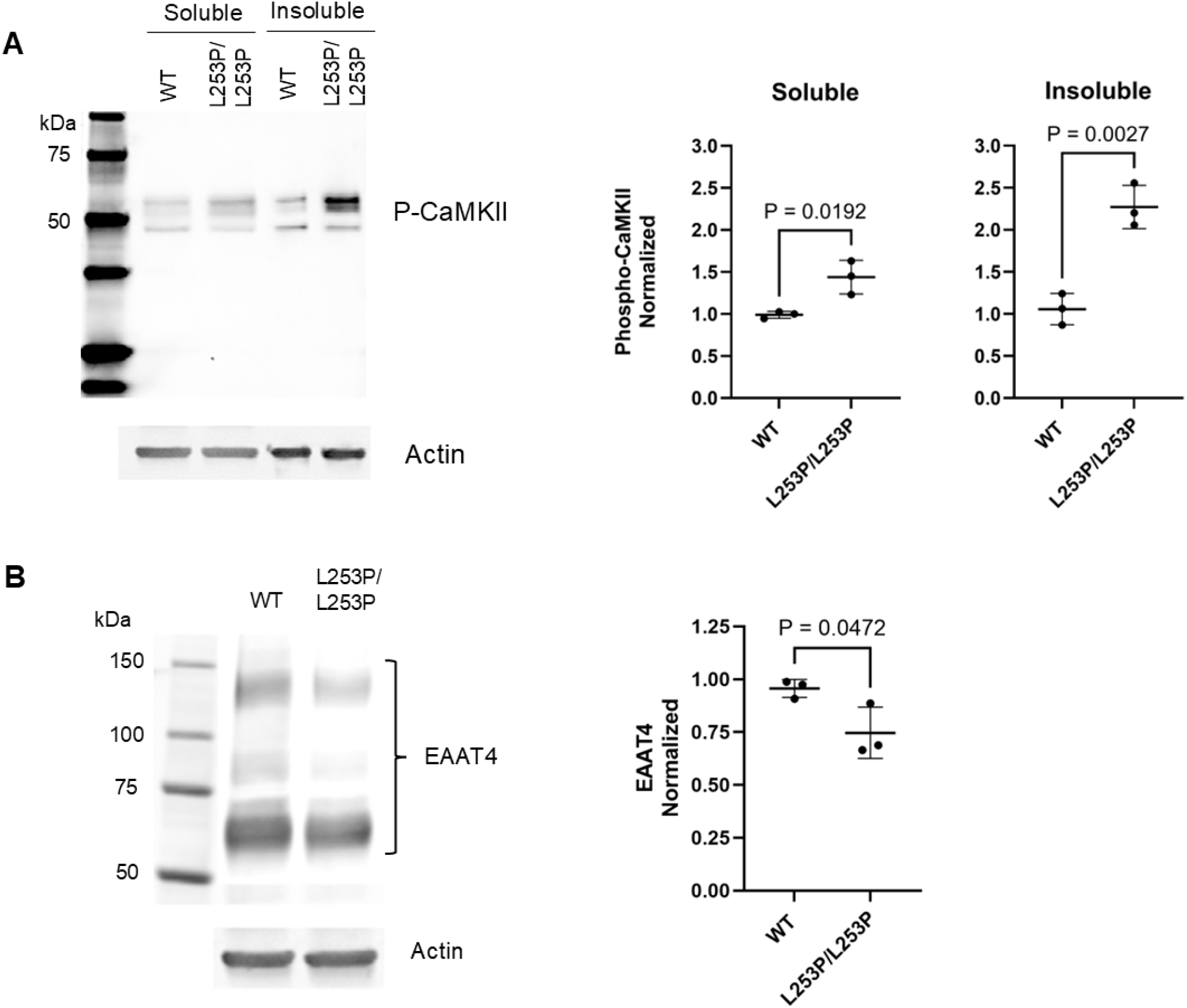
L253P activates CaMKII and decreases EAAT4 protein abundance. (A) Left, western blot of soluble and insoluble cerebellum protein fractions from wild-type and homozygous mice. Extracts were probed with a phospho-specific CaMKII antibody that recognizes active state of multiple CaMKII isoforms. Right, quantitation of P-CaMKII indicating increased activation of CaMKII in homozygous mice extracts relative to WT. (B) Left, western blot showing decreased EAAT4 protein in the soluble fraction of homozygous vs WT extracts. Right, quantitation showing reduced EAAT4 in homozygous cerebellum extracts relative to WT. Unpaired t-test, n=3.

Prior work in *β-III-spectrin* knockout mice showed that loss of β-III-spectrin results in reduced abundance the glutamate transporter EAAT4 (12, 15), potentially leading to glutamate excitotoxicity. Notably, EAAT4 was not identified as a β-III-spectrin interactor in our IP-MS experiments despite prior work supporting a physical interaction between the two proteins (21). Regardless, given the significance of EAAT4 to regulating glutamate signalling, we tested whether EAAT4 protein abundance is altered. Significantly, EAAT4 protein level was reduced by ~25% in *β-III-spectrin*^*L253P/L253P*^ tissue relative to WT, **Fig. 7B**. Consistent with this, in stained cerebellum slices, the EAAT4 fluorescence signal in Purkinje neurons appears weaker in *β-III-spectrin*^*L253P/L253P*^ mice compared to WT, **Fig. S4**. Altogether, these data suggest that L253P disrupts post-synaptic signalling involving glutamate and calcium signalling pathways. Moreover, these disruptions to CaMKII activity and EAAT4 agree with findings in *β-III-spectrin* knockout mice, further validating the L253P mouse as a system to further explore how L253P disrupts β-III-spectrin and Purkinje neuron functions, and for testing candidate SCA5 therapeutics.

## Discussion

Here we characterized a SCA5 *β-III-spectrin* knock-in mouse for *in vivo* investigations of disease mechanisms associated with the L253P mutation and for future therapeutic discovery. The L253P mouse shows mild motor impairment revealed by an increase in foot slips on an elevated beam. Mild motor deficits were similarly observed for the transgenic SCA5 E532_M544del mouse (20). This is consistent with the late onset and mild ataxia documented for both mutations in patients (2, 27). In contrast, motor impairment is much more pronounced in two independently developed *β-III-spectrin* knockout mice (12, 15). Likewise, severe motor developmental delay and intellectual disability result from homozygous frameshift or nonsense mutations, in the condition known as spectrin-associated autosomal recessive cerebellar ataxia 1 (SPARCA1)/spinocerebellar ataxia, autosomal recessive 14 (SCAR14) (28–30). We showed in the *β-III-spectrin*^*L253P/L253P*^ mice that CaMKII activity and EAAT4 protein level are disrupted, mirroring findings in *β-III-spectrin* knockout mice. However, given the much milder motor phenotype of the *β-III-spectrin*^*L253P/L253P*^ mouse compared to the *β-III-spectrin* knockout mice, significant differences must exist in the underlying molecular processes driving pathogenesis in the two model systems. The reproducible foot slip phenotype at 20 weeks makes the *β-III-spectrin*^*L253P/L253P*^ mouse amenable for testing efficacy of candidate SCA5 therapeutics. Moreover, given that L253P is one of many ABD-localized SCA5 mutations that causes increased actin-binding affinity (6), preclinical findings using the *β-III-spectrin*^*L253P/L253P*^ mouse may be broadly generalizable.

A central finding of this study is the pronounced redistribution of β-III-spectrin from distal dendrites to the soma and proximal dendrites of Purkinje neurons, accompanied by the formation of intracellular inclusions. The inclusions contain F-actin and α-II-spectrin, indicating that mutant β-III-spectrin retains the capacity to engage cytoskeletal partners and is unlikely to represent misfolded or denatured protein. Instead, we suggest that inclusion formations are a downstream consequence of the high-affinity binding of β-III-spectrin to F-actin. We previously reported in transfected HEK293 cells that L253P β-III-spectrin prominently localizes to the cell cortex but also accumulates on F-actin-rich intracellular vesicles (14). We speculated that high-affinity actin binding drives internalization of the mutant protein. In the mouse, the observation that small inclusions are distributed near the plasma membrane while larger inclusions accumulate perinuclearly supports a model in which mutant spectrin complexes originate at the cortex and undergo retrograde transport toward the cell center, potentially via dynein–dynactin–mediated mechanisms. Prior work in Drosophila supports a functional interaction between the L253P mutation and dynein/dynactin mediated transport (31), which may present as inclusion formations in Purkinje neurons. The recovery of proteins involved in Golgi-to-ER trafficking and microtubule transport in the β-III-spectrin interactome (**Fig. S5**) is consistent with this possibility.

The selective presence of inclusions in Purkinje neurons, but not hippocampal or cortical neurons despite similar cortical accumulation of β-III-spectrin, suggests that cell-type–specific factors influence inclusion formation and/or vulnerability. In addition to inclusion formation, the redistribution of spectrin to proximal membranes is likely to alter membrane trafficking dynamics (32, 33). Because spectrin networks can modulate clathrin-mediated endocytosis (34, 35), excess spectrin at proximal membranes combined with depletion from distal dendrites may create spatially opposing effects on membrane turnover and receptor trafficking. Such spatial dysregulation could contribute to impaired synaptic signaling even in the absence of overt neurodegeneration. The *β-III-spectrin*^*L253P/L253P*^ knock-in mouse model will allow such a mechanism of SCA5 pathogenesis to be explored within Purkinje neurons, as well as hippocampal and cortical neurons.

We conducted a proteomic analysis that identified an extensive β-III-spectrin interactome enriched for postsynaptic signaling components, including CaMKII subunits, glutamate receptor machinery, SERCA2, and additional calcium-regulated signaling complexes. Potentially, β-III-spectrin recruits CaMKII to regulate ion channels or other membrane proteins in Purkinje neurons. Significantly, prior work showed that CaMKII forms a signaling complex with β-IV-spectrin to regulate sodium channels in neurons (36). These findings support a model in which β-III-spectrin functions as a scaffold coordinating multiple elements of postsynaptic calcium regulation. Consistent with this interpretation, we observed increased CaMKII autophosphorylation in *β-III-spectrin*^*L253P/L253P*^ cerebellum, indicating altered intracellular calcium signaling. Reduced EAAT4 abundance may contribute to elevated synaptic glutamate and increased cytoplasmic calcium; however, the interactome suggests that additional mechanisms are likely involved. For example, altered association with SERCA2 could impair calcium sequestration into the endoplasmic reticulum, while disruption of NMDA receptors or CASK-related signaling complexes could influence calcium entry or downstream signaling. Thus, L253P appears to perturb calcium homeostasis through multiple converging pathways rather than a single dominant mechanism. Importantly, although CaMKII activation and EAAT4 reduction mirror findings in knockout mice, the substantially milder phenotype of the *β-III-spectrin*^*L253P/L253P*^ model indicates that pathogenic mechanisms differ quantitatively or qualitatively between loss-of-function and dominant mutations. One possibility is that L253P produces a combination of partial functional loss in distal dendrites together with toxic gain-of-function effects associated with abnormal cortical accumulation and inclusion formation. This dual mechanism may better reflect human SCA5 pathology.

The relevance of calcium dysregulation to disease is supported by recent studies demonstrating that T-type calcium channel inhibitors improve phenotypes in β-III-spectrin knockout models. In contrast, the FDA approved anti-glutamatergic drug, riluzole, did not improve motor symptoms in the knockout mouse despite reduced levels of EAAT4 (19). The *β-III-spectrin*^*L253P/L253P*^ mouse therefore will provide an important complementary model for evaluating calcium-targeted therapies as well as small-molecule modulators designed to normalize spectrin– actin interactions. In addition, beyond calcium signaling, the β-III-spectrin interactome implicates additional pathways potentially relevant to disease, including actomyosin regulation and membrane trafficking processes. These findings highlight the possibility that SCA5 pathogenesis arises from disruption of multiple interconnected cellular systems governing synaptic organization and neuronal architecture (37). The *β-III-spectrin*^*L253P/L253P*^ mouse provides a valuable platform for defining these mechanisms and for testing therapeutic strategies aimed at restoring spectrin function and neuronal signaling.

## Experimental procedures

### Statistical analysis

Statistical analysis was performed in Prism Graphpad v10.6. The type of statistical test and n values are indicated in figure legends.

### Mice

University of Minnesota Institutional Animal Care and Use Committee approved all mouse protocols. All mice were housed and managed by Research Animal Resources under pathogen-free conditions in an Association for Assessment and Accreditation of Laboratory Animal Care International approved facility. The mice had ad libitum access to food and water. In all experiments, similar numbers of male and female mice were used. All mice were age-matched within experiments and littermate controls were used when possible.

### Generation of the SCA5 *β-III-spectrin*^*L253P* /+^ mouse

CRISPR-Cas9 homology-directed repair was used to introduce mutations. The guide sequence sgRNA ACTTGGCCTGACGAAGCTCC was combined with Cas9 protein to generate a targeting complex. This was also combined with a single stranded DNA oligonucleotide GCTTTCAATCTGGCTGAAAAGGAACTTGGCCTGACGAAGCCCCTAGATCCTGAAGATGTGAATGTAGACCAGCCCGACGAGAAGT (Integrated DNA Technologies). The Cas9 guide RNA complex plus DNA template were injected into mouse single cell embryos. Two nucleotide alterations were introduced to generate the L253P mutation and destroy the PAM site, simultaneously creating a BfaI enzyme digest site for genotyping. Complexing and design was performed in conjunction with University of Minnesota Genome Engineering Shared Resource. CRISPR complex injection into fertilized FVB/NJ (Jackson Laboratory #001800) embryos was performed by the University of Minnesota Mouse Genetics Laboratory.

### Genotyping

DNA was extracted from tissue via Wizard Genomic DNA Purification Kit (Promega A2361) for use in PCR. PCR cycling conditions: 94°C 1min x1, 94°C 10sec, 60°C 30sec x35; 10μM TGA GTC CCT GAA GAA GTG TAA TG and GCC ACG TAG GTG ATG ATA GAC primers (Integrated DNA Technologies) plus 2 ul DNA are used in 20 μL GoTaq G2 master mix (Promega M7823). 10 ul of the PCR reaction was digested with 10 units BfaI at 37°C for 1 hour and 10 ul of the sample was mock treated minus enzyme. Samples were run side by side on a 2% agarose get looking for an uncut 143bp band or cut bands at 85 and 58 bp. Mice with mutations were Sanger sequenced. Once the founder mice sequences were confirmed they were mated with subsequent generations back crossed to FVB/NJ mice. Restriction digest fragement length g enotyping was transitioned to endpoint genotyping. Probes /56-FAM/AC GAA GCC C/ZEN/C TAG ATC CTG AAG ATG for L253P and /5HEX/AC GAA GCT C/ZEN/C TGG ATC CTG AAG ATG for wild-type were combine wtih the above primers. Primer and probe concentrations were optimized to 0.2 μM.Primetime in Gen Expression master mix (Integrated DNA Technologies 1055772). Cycling conditions: 94°C 5min x1, 94°C 10sec, 60°C 30sec x 40. Endpoint genotyping was used going forward.

### Behavioral analyses

University of Minnesota Behavior core provided blinded independent behavioral analysis. A single technician recorded hind foot slips throughout all the trials. Mice were habituated to the testing room for at least 15 min prior to the start of testing on each day. All testing apparatuses were cleaned between each animal with 70% ethanol. Males were run first then females. Observations were video recorded. Mice were trained for three days on a medium square beam followed by various beam trials on the fourth day. Beam sizes in order of testing: 16 mm square; 19 mm round; 10 mm wide square; 10mm diameter round. All beams are 36 inches long and 19 inches above the surface of the table. Mice start on a pedestal with a bright light. There is a dark box 7.5 × 7.5 × 5.5 inches (LxWxH) at the other end of the beam. Mice that fell off were placed back at the starting position and allowed to run again. If a mouse fell off twice they were excluded from that round of testing. Each mouse was tested twice on each beam, and hind quarter foot slips were counted.

### Immunofluorescence

Mice were deeply anesthetized and transcardially perfused with 10% buffered formalin, brains were extracted then stored in 10% formalin overnight. Brains were sunk in 30% sucrose overnight then sagittal sections were mounted in Tissue-Tek® O.C.T. compound (Fisher 23-730-571). Cryostat: 40-micron slices were cut and stored in phosphate buffered saline (PBS). Samples were washed 3 times in PBS gentle rocking 5 minutes each, floated in 1% Triton PBS for 10 minutes, and blocked in 5% donkey serum 0.3% Triton X-100 PBS for 1hr. After blocking, the sections were incubated for 24 hours at 4°C in 2% donkey serum 0.3% Triton X-100 in PBS buffer with rabbit anti-Sptbn2 (1:1,000, Novus NB110-58346), mouse anti-Sptbn2 (1:1,000, Santa Cruz sc-515737) mouse ankyrin R clone N388A/10 (1-500, NeuroMAB 75-380), guinea pig anti-calbindin (1:1500, Synaptic Systems 214 004), F-Actin 488 (1:10,000 Phalloidin Abcam AB176753). The sections were washed 3 times in PBS and incubated for 24 hours in 2% donkey serum 0.3% Triton X-100 in PBS containing secondary antibody. Cross absorbed donkey secondaries we purchased from Jackson Immunoresearch: Alexa Fluor 405, 488, 594, and 647 and were used at 1:1000 in various combinations depending on the primary antibody species while minimizing crosstalk. (Jackson Immumo 715-605-150, 715-545-150, 711-545-1552, 706-475-148, 706-605-148, 706-575-148)

### Fluorescence confocal imaging

A Leica Stellaris 8 confocal inverted microscope was used to acquire images. Images were acquired by thresholding to maximal intensity of the strongest sample signal for each channel. Images were collected across all samples within a group using the same parameters. Each image was collected at 1024×1024 with 2 times line averaging at 8 bit. Pinhole was set to 1 and there was no digital magnification. Z stacks were collected using system optimized steps. Data is stored with all metadata in raw .lif file format. Imaris v10.2 software was used to analyze the images for inclusions. To quantify volume of inclusions in the Purkinje neuron soma, somatic masks were created by combining calbindin and β-III-spectrin signals to create a surface. Imaris used the cell surface data to identify the Purkinje neuron soma based on minimum volume and sphericity. Masked soma area was used to train the Imaris’ AI FIJI plugin. The AI identified inclusions using minimum inclusion volume and fluorescent intensity of homozygous mouse samples. Once able to identify inclusions all images were batch processed. All training and pipeline pathways are available with the image files. Look up table parameters (color and intensity) were adjusted for better visualization in this publication. Raw data was not adjusted for analysis.

### Immunoprecipitation for mass spectrometry

Frozen WT, *β-III-spectrin*^*L253P/+*^ and *β-III-spectrin*^*L253P/L253P*^ mutant mouse cerebella, ages 25.4 – 26.0 weeks, were cut into small pieces in 2 mL of IP buffer (20 mM HEPES pH 7.5, 130 mM NaCl, 2 mM EDTA pH 7.5, 0.2% Triton X-100, 0.2% IGEPAL CA-630, 1X PhosSTOP (Sigma-Aldrich), 1X cOmplete mini, EDTA-free protease inhibitor (Roche)) before transferring to a 1 mL Dounce homogenizer. Each sample was homogenized twice on ice for a duration of 2 minutes each time. The samples were centrifuged at 10,000 x g at 4°C for 10 minutes. After that, each supernatant was clarified by incubating for 1 hour with 50 µL Protein A agarose beads, previously equilibrated with wash buffer (20 mM HEPES pH 7.5, 2 mM EDTA pH 7.5, 130 mM NaCl). The clarified supernatants were collected after spinning down at 750 x g for 1 minute at 4°C. Bradford assay was performed to measure the total protein concentration. Each supernatant sample was divided into two aliquots, each containing 2 mg total protein, and incubated with either 7 µg anti-β-III-spectrin antibody (Novus Biologicals) or 7 µg normal Rabbit IgG (Santa Cruz Biotechnology) for 1-2 hours, with inversion, at 4 °C. Beta-III-spectrin and control antibodies were then isolated by addition of 50 µL freshly equilibrated Protein A agarose beads and rotating at 4°C overnight. The Protein A/antibody complexes were collected by spinning at 750 x g at 4 °C for 1 minute. Beads were washed three times with 500 µL wash buffer. The IP samples were eluted into 60 µL 2x Laemmli sample buffer, made from 4x Laemmli sample buffer, after brief vortex followed by 5 minutes incubation at 95 °C. The IP samples were collected after spinning at 750 x g for 1 minute at room temperature and were diluted to 1x with water before loading samples on SDS-PAGE gels. The IP samples were run 2 cm distance into the resolving gel and gels stained with Coomassie G-250. For each WT and *β-III-spectrin*^*L253P/L253P*^ IP samples, a single ~2 cm long gel slice was isolated that contained all supernatant proteins. The gel slices were destained and sent on dry ice to the Integrated Mass Spectrometry Unit at Michigan State University for mass spectrometry analyses.

### In-gel protein digestion for mass spectrometry

Gel slices were cut into ~0.5 mm cubes and destained twice for 1 hr in 400 µl 25 mM ammonium bicarbonate (AMBIC):50% ACN. Cubes were dehydrated with 200 µl ACN for 5 min before removing solvent and dried to completion in a speed vacuum centrifuge at 30 °C. Cubes were resuspended in 200 µl 40 mM AMBIC:10% ACN with 1 µg trypsin (Promega, no. V5280) and 500 ng Lys-C (Promega, no. V1671) and incubated at 37 °C overnight. Solvent was isolated and cubes were dehydrated twice with 50% ACN:5% formic acid. Combined solvents were dried to completion in a speed vacuum centrifuge at 30 °C and resuspended in 50 µl 2% ACN, 0.1% formic acid. Samples were centrifuged at 14k x G for 2 min to remove residual gel pieces. 30 µl was taken for MS analysis.

### Nanoscale liquid chromatography coupled to tandem mass spectrometry

Nanoscale liquid chromatography coupled MS/MS separations were performed with a Thermo Scientific Ultimate 3000 RSLCnano System. Peptides were desalted in-line using a 3 μm diameter bead, C18 Acclaim PepMap trap column (75 μm × 20 mm) with 2% ACN, 0.1% formic acid (FA) for 8.75 min with a flow rate of 2 μL/min at 40 °C. The trap column was then brought in line with a 2 μm diameter bead, C18 EASY-Spray column (75 μm × 250 mm) for analytical separation over 127.5 min with a flow rate of 350 nL/min at 40 °C. The mobile phase consisted of 0.1% FA (buffer A) and 0.1% FA in ACN (buffer B). The separation gradient was as follows: 8.75 min desalting, 98.75 min 4–40% B, 2 min 40–65% B, 3 min 65–95% B, 11 min 95% B, 1 min 95–4% B, 3 min 4% B. Five microliters of each sample was injected. Top 20 data-dependent mass spectrometric analysis was performed with a Q Exactive HF-X Hybrid Quadrupole-Orbitrap Mass Spectrometer. MS1 resolution was 60K at 200 m/z with a maximum injection time of 45 ms, AGC target of 3e6, and scan range of 300–1500 m/z. MS2 resolution was 30K at 200 m/z, with a maximum injection time of 54 ms, AGC target of 1e5, and isolation range of 1.3 m/z. HCD normalized collision energy was 28. Only ions with charge states from +2 to +6 were selected for fragmentation, and the dynamic exclusion was set to 30 s. The electrospray voltage was 1.9 kV at a 2.0 mm tip to inlet distance. The ion capillary temperature was 280 °C, and the RF level was 55.0. All other parameters were set as a default.

### Mass spectrometry protein identification and quantitation

RAW mass spectrometry files were identified using Thermo Scientific Proteome Discoverer software (version 2.5) and Sequest against the *Mus musculus* and reviewed *Oryctolagus cuniculus* Uniprot proteome databases (UP000000589, 63,321 unique sequences; UP000001811, 1.069 unique sequences) additionally including trypsin (P00761), Lys-C (Q7M135, Q02SZ7), and contaminants. Enzyme specificity was set to trypsin, allowing up to 2 missed cleavages with an MS1 tolerance of 10 ppm and a fragment tolerance of 0.02 Da. Oxidation (M), acetylation (protein N-term), and methionine loss (protein N-term) were set as dynamic modifications. Peptide and protein false discovery rates (FDR) were 1% with the threshold determined via decoy search using the Percolator algorithm. At least two peptide identifications were required per protein identification. Peptide confidence was set to “High”. Label-free quantitation was performed using the Precursor Ions Quantifier node. Unique and Razor peptides were considered for quantitation. Precursor abundance was based on the peak intensity. Protein abundance was normalized by the total peptide amount. Protein abundance ratios were calculated using the average values of normalized abundance replicates. All other parameters were set as default.

### STRING cluster analysis

The 157 proteins composing the β-III-spectrin interactome were analyzed in the STRING database for clusters. K-means cluster analysis was performed with number of clusters set to 10. Under Settings, meaning of cluster edges was set to “confidence,” such that increasing line thickness means greater confidence of an interaction between two proteins. All other settings left to default.

### Western blot analyses

Frozen WT, *β-III-spectrin*^*L253P/+*^ and *β-III-spectrin*^*L253P/L253P*^ mutant mouse cerebella, ages 25.4 – 26.0 weeks, were cut into small pieces in 2 mL of IP buffer (20 mM HEPES pH 7.5, 130 mM NaCl, 2 mM EDTA pH 7.5, 0.2% Triton X-100, 0.2% IGEPAL CA-630, 1X PhosSTOP (Sigma-Aldrich), 1X cOmplete mini, EDTA-free protease inhibitor (Roche)) before transferring to a 1 mL Dounce homogenizer. Each sample was homogenized twice on ice for a duration of 2 minutes each time. The samples were centrifuged at 10,000 x g at 4°C for 10 minutes. Bradford assay was performed to measure the total protein concentration in the lysate supernatant while the pellet was saved on ice. After normalizing the concentrations of the proteins to the lowest concentration, soluble samples were made by mixing equal amounts of the sample with 2x Laemmli sample buffer (BioRad). To obtain comparable insoluble samples, 4x Laemmli sample buffer was added onto the pellet at volume equal to one tenth of the normalized supernatant total volume. The pellets were boiled at 95°C for 5 minutes followed by brief vortex. The insoluble samples were collected after centrifugation at 10,000 x g for 5 minutes at room temperature. Soluble and insoluble samples were resolved by SDS-PAGE gel followed by transferring to Immoblin-FL membranes (Millipore). β-III-spectrin proteins were detected by probing with anti-β-III-spectrin antibody (1:100; Santa Cruz A-8) diluted in 1% casein/1x PBS and 0.1% Tween-20 and IRDye 800CW secondary antibody (1:5000; LICORbio) diluted in 0.1% Tween-20, 0.01% SDS, 1% casein, 1x PBS. The membranes were imaged using Azure Sapphire 800 nm channel. Band intensity analyses were performed using Image Studio Lite (LICORbio). *β-III-spectrin*^*L253P/+*^ and *β-III-spectrin*^*L253P/L253P*^ soluble and insoluble β-III-spectrin signals were normalized to WT β-III-spectrin soluble and insoluble signals, respectively, loaded onto the same gels.

## Supporting Information

This article contains supporting information.

## Acknowledgments

We thank Dr. Robyn T. Rebbeck for critical reading of the manuscript.

## Author contributions

T.S.H., H.T.O, and A.W.A conceived and designed experiments. B.L.O., S.A.D., and M.T.T. performed experiments. A.W.A., M.T.T., E.M.C., J.L., and D.G.O. analyzed data. A.W.A wrote the manuscript. T.S.H and B.L.O critically read and revised the manuscript.

## Funding and additional information

This work was supported by the NIH grants 1R35NS127248-01 to H.T.O., GM044757, R61NS11075, R33NS11075 to T.S.H, and 2R15NS116511-02 and a National Ataxia Foundation Early Career award to A.W.A.

## Conflict of interest

The authors declare no conflict of interest.

## Notes

### Competing Interest Statement

The authors have declared no competing interest.

