## Supplemental Information for "Impaired motor activity in a CRISPR SCA5 L253P knock-in mouse is associated with selective β-III-spectrin subcellular redistribution in the cerebellum"

**Fig. S1. L253P does not impact rotarod performance at 6 and 24 weeks.**

**Fig. S1.**

**A**

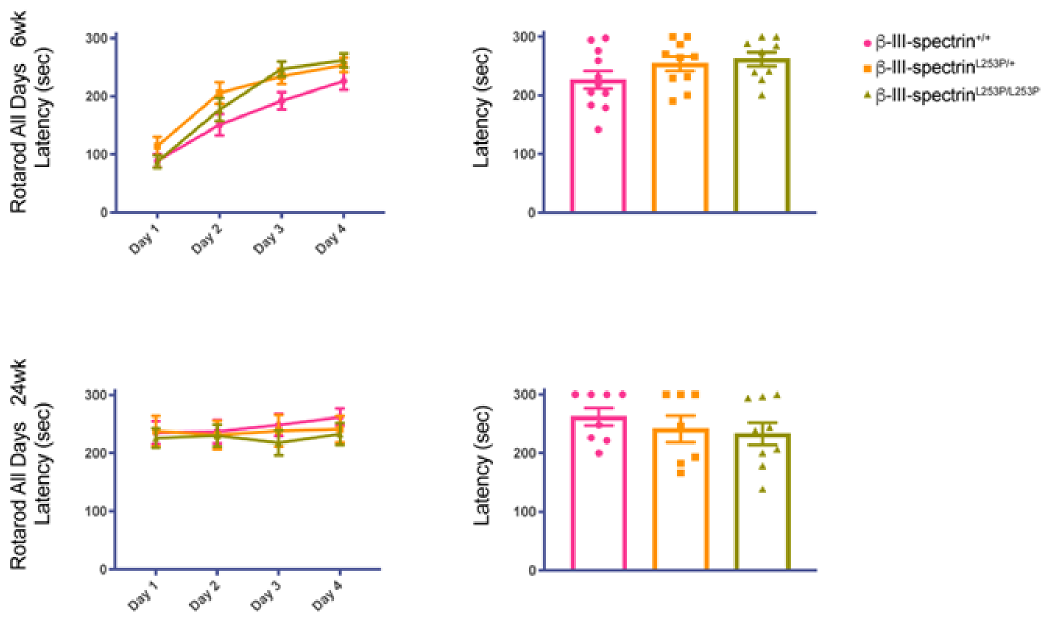

**Fig. S2. Detection of β-III-spectrin in Purkinje neurons using an antibody targeting the N-terminus of β-III-spectrin.**

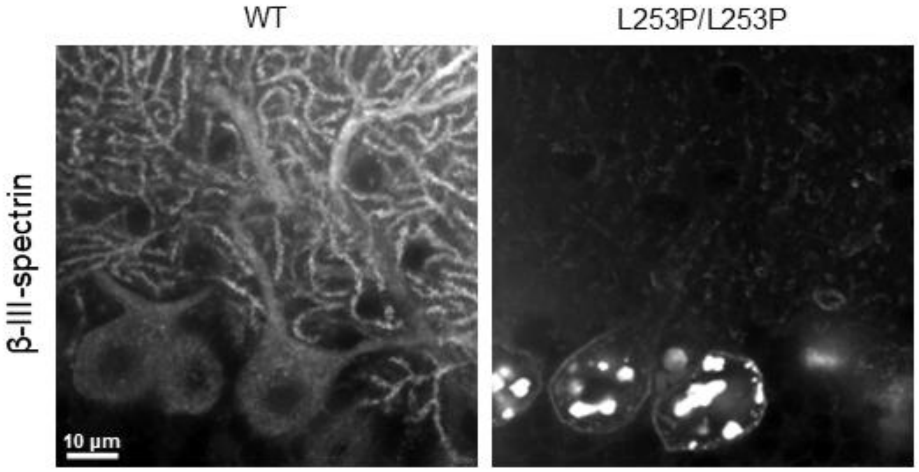

**Fig. S3. Small β-III-spectrin inclusions localizing near plasma membrane.**

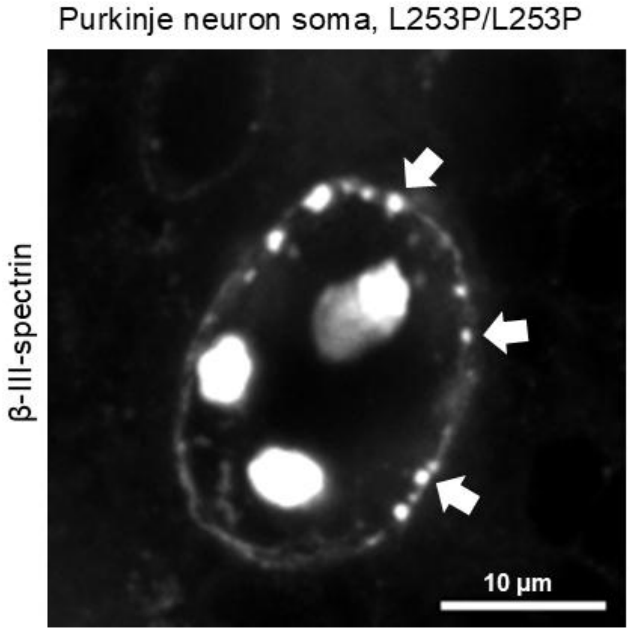

**Fig. S4. Neither ankyrin-R nor EAAT4 localize to β-III-spectrin inclusions.**

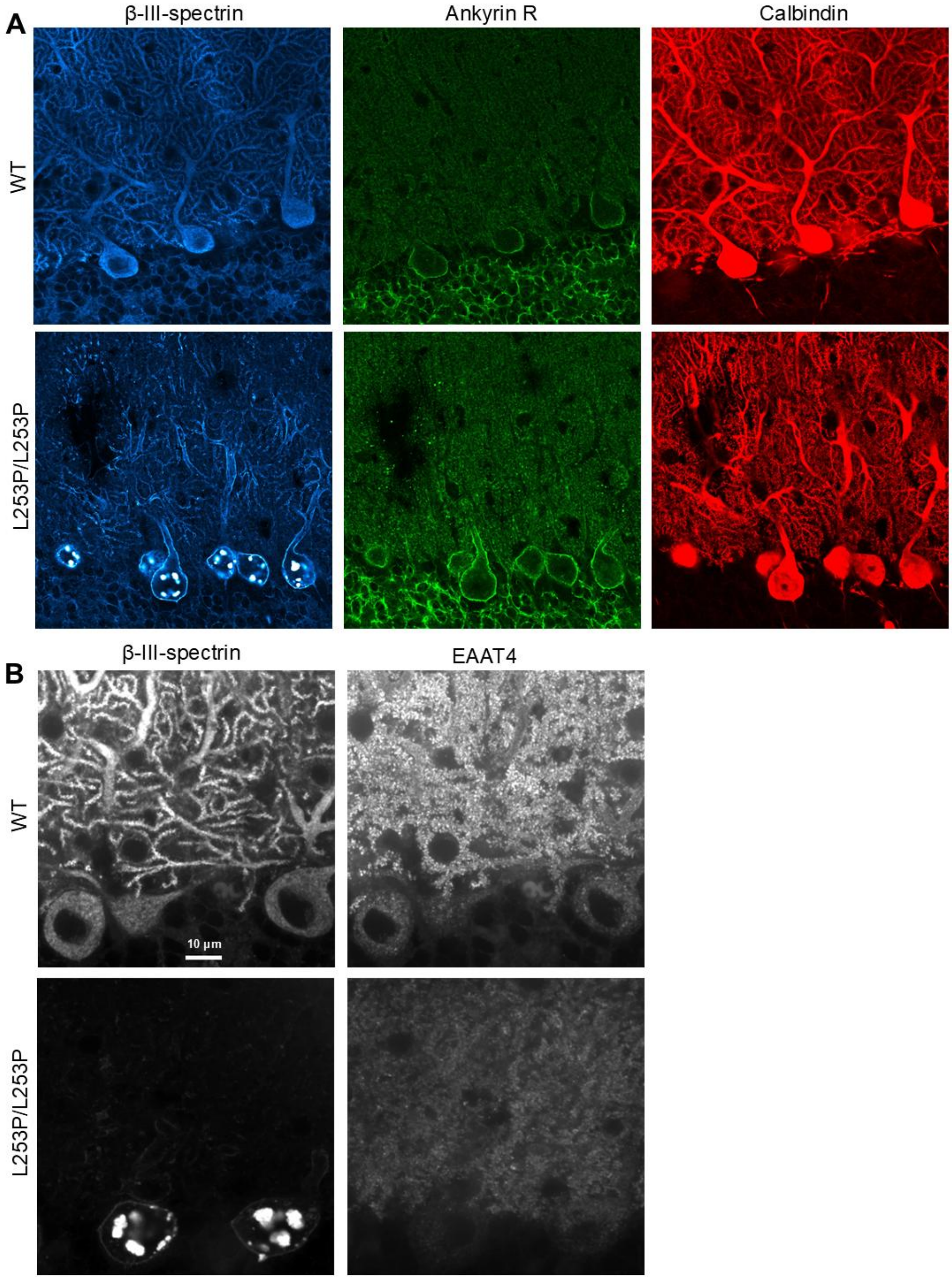

**Fig. S5. A cerebellum-specific β-III-spectrin protein interactome.**

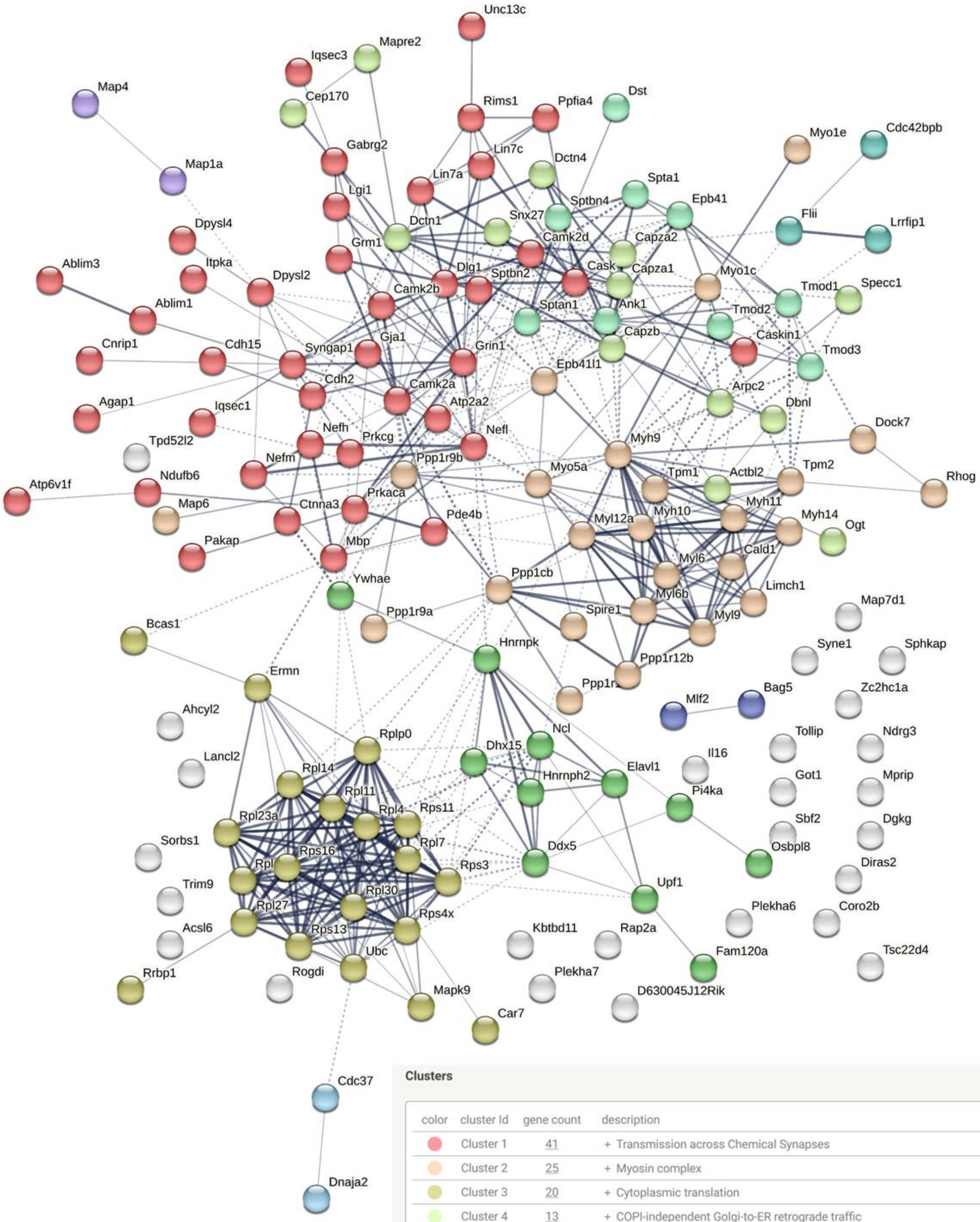


**Clusters**

| color | cluster Id | gene count | description |
|---|---|---|---|
| | Cluster 1 | 41 | + Transmission across Chemical Synapses |
| | Cluster 2 | 25 | + Myosin complex |
| | Cluster 3 | 20 | + Cytoplasmic translation |
| | Cluster 4 | 13 | + COPI-independent Golgi-to-ER retrograde traffic |
| | Cluster 5 | 11 | RNA recognition motif domain, and U2-type post-mRNA release spliceosomal comp... |
| | Cluster 6 | 9 | + Actin filament capping |
| | Cluster 7 | 3 | Cdc42bpb, Flii, Lrrfip1 |
| | Cluster 8 | 2 | Regulation of HSF1-mediated heat shock response, and HSP40/DnaJ peptide-bindi... |
| | Cluster 9 | 2 | Bag5, Mlf2 |
| | Cluster 10 | 2 | Map1a, Map4 |

### Figure legends

**Fig. S1. L253P does not impact rotarod performance at 6 and 24 weeks.** Two naïve cohorts were assessed. N values for 6 week cohort: WT, 11; L253P/+, 10; L253P/L253P, 9. N values for 24 week cohort: WT, 8; L253P/+, 7; L253P/L253P, 9.

**Fig. S2. Detection of β-III-spectrin in Purkinje neurons using an antibody targeting the N-terminus of β-III-spectrin.** Representative confocal images of 20 week mouse Purkinje neurons.

**Fig. S3. Small β-III-spectrin inclusions localizing near plasma membrane.** Representative 63x confocal image from 20 week homozygous mouse. White arrows indicate small inclusions near plasma membrane.

**Fig. S4. Neither ankyrin-R nor EAAT4 localize to β-III-spectrin inclusions.** Representative 63x confocal images from WT and homozygous mice.

**Fig. S5. A cerebellum-specific β-III-spectrin protein interactome.** The interactome contains 157 proteins. Cluster analysis was performed using the STRING database.
